# Sex determining gene transposition as an evolutionary platform for chromosome turnover

**DOI:** 10.1101/2020.03.14.991026

**Authors:** Fernando Ayllon, Monica Favnebøe Solberg, François Besnier, Per Gunnar Fjelldal, Tom Johnny Hansen, Anna Wargelius, Rolf Brudvik Edvardsen, Kevin Alan Glover

## Abstract

Despite the key role that sex-determination plays in evolutionary processes, it is still poorly understood in many species. In salmonids, which are the best studied family of fishes, the master sex-determining gene sexually dimorphic on the Y-chromosome (*sdY*) has been identified. However, *sdY* displays unexplained discordance to the phenotypic sex, with a variable frequency of phenotypic females being reported as genetic males. Multiple sex determining loci in Atlantic salmon have also been reported, possibly as a result of transposition, suggesting a recent and non-random sex chromosome turnover in this species. We hypothesized the existence of an autosomic pseudocopy of *sdY* that is transmitted in accordance with Mendelian inheritance. To test this we developed a qPCR methodology to detect the number of *sdY* copies present in the genome. Based on the observed phenotype/genotype frequencies and linkage analysis among 2025 offspring from 64 pedigree-controlled families of accurately phenotyped Atlantic salmon, we identified both males and females carrying one or two autosomic copies in addition to the Y-specific copy present in males. Copy number frequencies were consistent with Mendelian inheritance. Pseudocopy loci were mapped to different chromosomes evidencing non-random transitions of the sex determining gene in Atlantic salmon and the existence of functional constraints for chromosome turnover.

## Introduction

Most eukaryotic organisms reproduce sexually, yet the nature of the sexual system and the mechanism of sex determination often vary remarkably, even among closely related species (Ashman et al. 2014; Pennell, Mank, and Peichel 2018). This is in particularly true for teleosts where some species display genetic sex determination, some display environmental sex determination, while others a mixture of both (Heule, Salzburger, and Bohne 2014). Furthermore, both male- (XX females and XY males) and female-heterogametic systems (ZZ males and ZW females) are found evenly in closely related species such as tilapias (Cnaani et al. 2008) and sticklebacks (Ross et al. 2009).

Atlantic salmon (*Salmo salar*) is an anadromous fish inhabiting temperate streams in the north Atlantic. It belongs to the family Salmonidae, which includes multiple species from 11 genera including salmon, trout, charr, freshwater whitefishes, ciscoes and graylings. Globally, Atlantic salmon represents one of the most economically significant and iconic species, providing extensive angling recreation, extensive aquaculture production, and symbolizing healthy ecosystems in the rivers it inhabits. As a consequence, it is also one of the most exhaustively studied fish. The Atlantic salmon’s ancestor underwent a whole-genome duplication event approximately 88-103 million years ago (Macqueen and Johnston 2014), and is now in the process of rediploidization. As a result of this process, the Atlantic salmon genome consists of many paralogous regions (Lien et al. 2016) which in principle can diversify (Kjaerner-Semb et al. 2016) and acquire new functions as has been observed in other species displaying duplicated genomes (Qian and Zhang 2014). Interestingly, the presence of transposable elements found in the genome is among the highest found in vertebrates (Lien et al. 2016).

Genetic sex determination can be polymorphic (Bulmer and Bull 1982; Liew et al. 2012; Moore and Roberts 2013) but it may also be controlled by a single master sex determining gene (Meise et al. 1998; Pan et al. 2019). Such genes are likely to function by initiating the developmental cascade which ends in the development of ovaries or testes. In fish, a number of master sex determining (MSD) genes have been identified, most of which belong to one of three protein families (DMRT, SOX, and TGF-ß and their signaling pathways). The Salmonidae is an exception to this, where the MSD is *sexually dimorphic on the Y-chromsome* (*sdY*) first discovered in rainbow trout (*Oncorhynchus mykiss*) (Yano et al. 2012; Yano et al. 2013). The immune related factor *sdY* blocks the female differentiation pathway by interacting with the transcription factor Foxl2, thereby allowing male differentiation (Bertho et al. 2018). The discovery of *sdY*, and the subsequent development of assays for rapid genetic sex determination has opened novel possibilities. For example, molecular assays have been used to determine genetic sex in adults that were not phenotyped but subsequently used for sex-specific studies such as investigation into the genetic basis of age at maturity (Ayllon et al. 2015; Barson et al. 2015; Ayllon et al. 2019).

Several studies have reported a discordance between phenotypic and *sdY* sex within the Salmonidae family (Eisbrenner et al. 2014; Cavileer et al. 2015; Larson et al. 2016; Podlesnykh, Brykov, and Kukhlevsky 2017). Discordance between genetic and phenotypic sex is not uncommon in fishes, and it is typical in a species displaying a combination of genetic and environmental sex determination (Hattori et al. 2019). However, environmental sex determination has not been reported in the family Salmonidae, and several alternative theories for this discordance have been put forward (Yano et al. 2012). Nevertheless, the mechanisms underpinning this discordance are unknown. Adding to the complexity of the situation is the fact that *sdY* has been mapped to different regions of the genome in the various salmonid species, but also within the same species, suggesting that it transposes to a new location either at the time of speciation (Phillips 2013) or more recently (Kijas et al. 2018). Specifically within Atlantic salmon, *sdY* has been mapped to chromosomes Ssa02, Ssa03, Ssa06 and possibly Ssa21 (Eisbrenner et al. 2014; Lubieniecki et al. 2015; Kijas et al. 2018; Gabian et al. 2019).

In this study we determined why genetic sex determination does not always correlate with phenotypic sex in Atlantic salmon. We first asked the question whether the observed discordance could be linked to non-functional copies of *sdY* locus in the genome. We thereafter answered this by quantifying multiple *sdY* copies in the genome by performing qPCR on genomic DNA from in 2025 accurately phenotyped Atlantic salmon originating from 64 families of domesticated, F1-hybrid, and wild origin. We demonstrate that the *sdY* gene has an infrequent pseudocopy in the genome, that is inherited in an autosomic mendelian fashion, which explains the observed discordance.

## Methods

### Experimental crosses

Over the past decade, we have studied an experimental population of domesticated and wild Atlantic salmon and their crosses at the aquaculture facility owned by the Institute of Marine Research located in Matre, western Norway (Solberg et al. 2013; Solberg et al. 2014; Ayllon et al. 2015; Harvey et al. 2016; Glover et al. 2018; Harvey et al. 2018). The reader is directed to these publications for full details regarding the standard rearing conditions experienced in this fish farm. In the present study, we produced a total of 29 (F1-C2011) and 39 (F1-C2012) experimental families in the years 2011 and 2012 respectively. These families originated from the domesticated Mowi strain (13 families), the wild Figgjo population (14 families), reciprocal F1-hybrids between Mowi and Figgjo (24 families), the wild Vosso population (7 families), and the wild Arna population (6 families). Full details of these experimental crosses and the background of the source populations are available elsewhere (Solberg et al. 2014).

After fertilization in 2010 and 2011, eggs were incubated in single-family containers until the eyed stage when they were mixed into common-garden experiments to study a range of phenotypic traits (data not used here). These fish were first reared until smoltification in freshwater aged 1+ when 2000 (F1-C2011) and 2400 (F1-2012) individuals were PIT tagged and DNA sampled, and thereafter transferred into sea-cages where they were reared until they matured after a further 1-3 years. Families represented by less than 10 individuals at maturity were discarded. Upon maturation, the phenotypic sex of 2025 individuals from 64 families was accurately recorded by dissection, giving a total of 1048 and 977 phenotypic males and females respectively.

### Genetic analysis – microsatellites and SNPs

Total DNA from all offspring and parents was purified using the Qiagen DNeasy Blood & Tissue Kit (Qiagen, Hilden, Germany) according to the manufacturer’s recommendations Microsatellite DNA parentage testing was used to unambiguously identify the pedigree of all individuals used in this study using the exclusion based method implemented in FAP (Taggart 2007). The laboratory conducting these analyses has extensive experience in DNA parentage testing (Solberg et al. 2013; Solberg et al. 2014; Harvey et al. 2016; Glover et al. 2018; Harvey et al. 2018), and the full details regarding the markers used and their amplification conditions are available in these previous studies. In addition to microsatellites, a set of 116 genome-wide distributed SNPs were genotyped in all offspring and parents for the purpose of linkage mapping (see below). This analysis was performed on a MassARRAY Analyzer 4 from Agena Bioscience™ according to the manufacturer’s instructions. The final dataset for mapping included 109 genome-wide distributed SNPs once those displaying poor coverage and clustering were removed. The list of SNPs and their positions area available elsewhere (Besnier et al. 2020).

### PCR-based *sdY* tests

The *sdY* presence / absence was validated by a PCR-based methodology aimed to detect the presence of the *sdY* gene (Yano et al. 2012; Eisbrenner et al. 2014). Individuals showing amplicons of exon 2 and 4 were designated as males. As a positive PCR control and for species determination we used the presence of the 5S rRNA gene (Pendas et al. 1995). PCR amplifications were performed using reaction mixtures containing approximately 50 ng of extracted Atlantic salmon DNA, 10 nM Tris–HCl pH 8.8, 1.5 mM MgCl_2_, 50 mM KCl, 0.1% Triton X-100, 0.35 μM of each primers, 0.5 Units of DNA Taq Polymerase (Promega, Madison, WI, USA) and 250 μM of each dNTP in a final volume of 20 μL. PCR products were visualized in 3% agarose gels.

A quantitative PCR (qPCR) based methodology was developed to quantify the number of *sdY*-liked copies present. *gapdh*, *sdY* exon 2 and *sdY* exon4 were multiplexed using 5’labelled probes (Suppl. Table 1). The *gapdh* locus was used as an internal positive control (IPC) and reference gene to estimate fold change (FC) values (Livak and Schmittgen 2001). Amplification reactions were run on a Quanstudio5 384-system real time detection system (Thermo Fisher Scientific, USA). Reactions consisted of a Pre-Read stage (60°C for 30s), a Hold Stage (95°C for 10min), a PCR stage (40 cycles of 95°C for 15s and 60°C for 1min) and Post-Read stage (60°C for 30s). Each 5 μl reaction contained the following final concentrations: 1x Taqman Universal MasterMix, 1μM *gapdh* forward and reverse primers, 0.2 μM *gapdh* TaqMan probe, 1.4μM sdy_Exon2 forward and reverse primers, 0.32 μM sdy_Exon2 TaqMan probe, 2.1 μM sdy_Exon4 forward and reverse primers, 0.48 μM sdy_Exon4 TaqMan probe and 2 ng/μl of gDNA template. For every 384 well reaction plate we included 8 to 16 reference males and females and a minimum of 8 no template controls (NTC).

**Table 1.** Overview of the 64 families including: family unique identifier (ID), strain origin (Type), Dam and sire unique identifier (Dam ID and Sire ID respectively), total number of individuals per family (N) and Dam and sire *sdY* copy number (Dam *sdY* and Sire *sdY* respectively). Additionally, linkage analysis results for the actual sex determining *sdY* locus (SEX) and the pseudocopy with *P* values. NA values are provided when analysis could not be performed either due to low N or insufficient genetic variation.

| Family |  |  |  |  |  |  | SEX Linkage Analysis |  | *sdY* Pseudocopy Linkage Analysis |  |
| --- | --- | --- | --- | --- | --- | --- | --- | --- | --- | --- |
| ID | Type | Dam ID | Sire ID | N | *sdY* Dam | *sdY* Sire | Chromosome | *P* value | Chromosome | *P* value |
| F01 | Farm | #01 | #11 | 27 | 0x | 1x | Ssa02 | 2E-10 | NA | NA |
| F02 | Hybrid | #01 | #31 | 70 | 0x | 1x | Ssa03 | 5E-08 | NA | NA |
| F03 | Farm | #02 | #12 | 25 | 0x | 1x | NA | NA | NA | NA |
| F04 | Hybrid | #02 | #32 | 37 | 0x | 1x | NA | NA | NA | NA |
| F06 | Hybrid | #03 | #32 | 23 | 0x | 1x | NA | NA | NA | NA |
| F07 | Farm | #04 | #14 | 22 | 0x | 1x | Ssa02 | 2E-20 | NA | NA |
| F08 | Hybrid | #04 | #33 | 27 | 0x | 1x | Ssa06 | 4E-10 | NA | NA |
| F09 | Farm | #05 | #15 | 36 | 0x | 1x | Ssa02 | 2E-15 | NA | NA |
| F10 | Hybrid | #05 | #33 | 50 | 0x | 1x | Ssa06 | 4E-15 | NA | NA |
| F11 | Farm | #06 | #16 | 22 | 0x | 1x | Ssa02 | 3E-20 | NA | NA |
| F12 | Hybrid | #06 | #34 | 39 | 0x | 2x | Ssa02/Ssa03 | 2E-06/3E-06 | Ssa03 | 6E-05 |
| F13 | Farm | #07 | #17 | 29 | 0x | 1x | Ssa02 | 7E-09 | NA | NA |
| F14 | Hybrid | #07 | #35 | 50 | 0x | 2x | Ssa02 | 8E-05 | Ssa06 | 3E-4 |
| F16 | Hybrid | #08 | #36 | 64 | 0x | 1x | Ssa03 | 3E-16 | NA | NA |
| F17 | Farm | #09 | #19 | 46 | 0x | 1x | Ssa02 | 2E-11 | NA | NA |
| F18 | Hybrid | #09 | #37 | 44 | 0x | 1x | Ssa06 | 2E-05 | NA | NA |
| F19 | Farm | #10 | #20 | 27 | 0x | 1x | Ssa02 | 7E-14 | NA | NA |
| F20 | Hybrid | #10 | #38 | 42 | 0x | 1x | NA | NA | NA | NA |
| F21 | Wild | #21 | #31 | 25 | 0x | 1x | Ssa03 | 1E-02 | NA | NA |
| F22 | Wild | #22 | #32 | 34 | 0x | 1x | Ssa06 | 1E-02 | NA | NA |
| F23 | Wild | #23 | #33 | 36 | 0x | 1x | Ssa06 | 4E-10 | NA | NA |
| F26 | Wild | #26 | #35 | 24 | 1x | 2x | Ssa02 | 1E-02 | Ssa28 | 1E-03 |
| F27 | Wild | #27 | #35 | 28 | 1x | 2x | Ssa02 | 2E-17 | Ssa06 | 3E-02 |
| F29 | Wild | #29 | #37 | 20 | 0x | 1x | Ssa06 | 2E-04 | NA | NA |
| F30 | Wild | #30 | #38 | 24 | 0x | 1x | Ssa06 | 2E-03 | NA | NA |
| K01 | Wild | A01 | A05 | 23 | 0x | 1x | Ssa02 | 1E-02 | NA | NA |
| K02 | Wild | A01 | A06 | 19 | 0x | 1x | Ssa06 | 6E-05 | NA | NA |
| K03 | Wild | A02 | A07 | 24 | 0x | 1x | Ssa06 | 2E-16 | NA | NA |
| K04 | Wild | A02 | A08 | 27 | 0x | 1x | Ssa03 | 1E-04 | NA | NA |
| K05 | Wild | A03 | A09 | 26 | 0x | 1x | Ssa02 | 2E-14 | NA | NA |
| K06 | Wild | A04 | A10 | 30 | 0x | 1x | Ssa06 | 2E-13 | NA | NA |
| K15 | Wild | V01 | V09 | 41 | 0x | 1x | Ssa02 | 1E-13 | NA | NA |
| K16 | Wild | V02 | V10 | 11 | 0x | 1x | Ssa02 | 3E-15 | NA | NA |
| K17 | Wild | V03 | V11 | 29 | 0x | 1x | Ssa06 | 4E-06 | NA | NA |
| K18 | Wild | V04 | V12 | 42 | 0x | 1x | NA | NA | NA | NA |
| K19 | Wild | V05 | V13 | 42 | 0x | 2x | NA | NA | NA | NA |
| K20 | Wild | V06 | V14 | 28 | 0x | 1x | Ssa02 | 2E-15 | NA | NA |
| K22 | Wild | V08 | V16 | 30 | 1x | 1x | Ssa02 | 1E-02 | Ssa06 | 6E-03 |
| K23 | Wild | F01 | F10 | 29 | 0x | 1x | Ssa06 | 7E-04 | NA | NA |
| K24 | Hybrid | F01 | M09 | 36 | 0x | 1x | Ssa06 | 1E-08 | NA | NA |
| K25 | Hybrid | M01 | F10 | 12 | 0x | 1x | Ssa06 | 2E-08 | NA | NA |
| K29 | Hybrid | M2 | F12 | 31 | 0x | 1x | Ssa06 | 3E-07 | NA | NA |
| K31 | Wild | F03 | F13 | 37 | 0x | 1x | Ssa02/Ssa06 | 2E-07/3E-06 | NA | NA |
| K32 | Hybrid | F03 | M11 | 28 | 0x | 1x | Ssa02 | 2E-14 | NA | NA |
| K33 | Hybrid | M3 | F13 | 32 | 0x | 1x | Ssa02 | 3E-30 | NA | NA |
| K34 | Farm | M03 | M11 | 26 | 0x | 1x | Ssa02 | 5E-25 | NA | NA |
| K37 | Hybrid | M04 | F14 | 52 | 0x | 1x | Ssa06 | 3E-19 | NA | NA |
| K38 | Farm | M04 | M12 | 15 | 0x | 1x | Ssa02 | 2E-10 | NA | NA |
| K39 | Wild | F05 | F15 | 22 | 0x | 1x | Ssa02 | 8E-06 | NA | NA |
| K40 | Hybrid | F05 | M13 | 31 | 0x | 1x | Ssa02 | 7E-12 | NA | NA |
| K41 | Hybrid | M05 | F15 | 12 | 0x | 1x | Ssa02 | 2E-08 | NA | NA |
| K43 | Wild | F06 | F16 | 26 | 0x | 1x | NA | NA | NA | NA |
| K44 | Hybrid | F06 | M14 | 29 | 0x | 1x | Ssa03 | 2E-04 | NA | NA |
| K45 | Hybrid | M6 | F16 | 39 | 0x | 1x | NA | NA | NA | NA |
| K46 | Farm | M6 | M14 | 21 | 0x | 1x | Ssa03 | 2E-09 | NA | NA |
| K47 | Wild | F07 | F17 | 38 | 0x | 1x | Ssa02 | 4E-25 | NA | NA |
| K48 | Hybrid | F07 | M15 | 48 | 0x | 1x | NA | NA | NA | NA |
| K49 | Hybrid | M7 | F17 | 53 | 0x | 1x | Ssa02 | 2E-30 | NA | NA |
| K50 | Farm | M07 | M15 | 17 | 0x | 1x | Ssa06 | 2E-02 | NA | NA |
| K51 | Wild | F08 | F18 | 39 | 0x | 2x | Ssa02 | 2E-04 | Ssa05 | 1E-03 |
| K52 | Hybrid | F08 | M16 | 54 | 0x | 1x | Ssa02 | 8E-13 | NA | NA |
| K53 | Hybrid | M08 | F18 | 31 | 0x | 2x | Ssa02 | 3E-04 | Ssa13 | 2E-03 |
| K54 | Farm | M8 | M16 | 22 | 0x | 1x | Ssa02 | 3E-04 | NA | NA |
| K55 | Wild | F09 | F11 | 15 | 0x | 1x | NA | NA | NA | NA |

In order to validate the qPCR methodology we used XY males and YY super-males. XY males will carry a single copy of the Y-specific sex determining gene *sdY*. On the other hand, super males will carry two *sdY* gene copies, one per Y chromosome. YY males are the product of either self-fertilization or double haploid males. Full details on YY super-males production can be found in Fjelldal et al. (*in prep.*). Briefly, eggs and milt from a hermaphrodite salmon were surgically removed to prevent undesired self-fertilization. Eggs were then self-fertilized either with normal or UV-irradiated milt. Following the fertilization with UV-irradiated milt, pressure mediated diploidization was carried out to produce the double haploid males used in this study.

### Linkage mapping

Linkage mapping was performed on all 64 families, including the 8 families showing discrepancy between genetic and phenotypic sex. For each family, the coefficient of Identity By Descent (IBD) among offspring alleles was estimated from both pedigree and genotype information as in (Pong-Wong et al. 2001). First, the genomic location of the sex determining locus was considered. Association between the binary phenotype (male / female) and the two paternally (maternally) inherited alleles was investigated by fitting a Chi-squared test in each family separately, at each SNP locus. Second, the genomic location of the sex discrepancy for the eight affected families was investigated following the same Chi-squared approach. Here, the two phenotypes were no longer male and female but discrepant / non-discrepant individuals, where the non-discrepant group consisted of all the regular males and females, and the discrepant group consisted of phenotypic females that amplified one or more copy of *sdY*, as well as phenotypic males that amplified more than one *sdY* copy.

### Statistical Analysis

Chi square tests with computed p values by Monte Carlo simulations (10^6^ replicates) were used to test for deviations of the observed values from the expected frequencies. All statistical analyses were conducted in R V.3.6.2. (R-Development-Core-Team 2019).

### Ethical considerations and research permits

The experimental protocols (permit numbers 4268, 5296) were approved by the Norwegian Animal Research Authority (NARA). Use of experimental animals were performed in strict accordance with the Norwegian Animal Welfare Act. This included anesthesia or euthanasia of fish using metacain (Finquel^®^ Vet, ScanVacc, Årnes, Norway), during all described procedures. In addition, all personnel involved in this experiment had undergone training approved by the Norwegian Food Safety Authority, which is mandatory for all personnel running experiments involving animals included in the Animal Welfare Act.

## Results

We found PCR-based discordance between the validated phenotypic sex and *sdY* genotype in 66 individuals across 64 families and 2025 fish (Fig. 1). All of the reported cases were phenotypic females displaying a positive signal for *sdY*. Discordance between phenotypic and genetic sex was only observed in 8 of the 64 families, ranging from 36 to 82% discordance among females per family. Of the 88 parents used as broodstock, three phenotypic females were *sdY* positive. These females were the parents of families F26, F27 and K22, all of which had offspring displaying discordance between phenotypic and genetic sex (Fig. 1). The five other families containing discordant offspring did not have discordant mothers.

**Figure 1.**
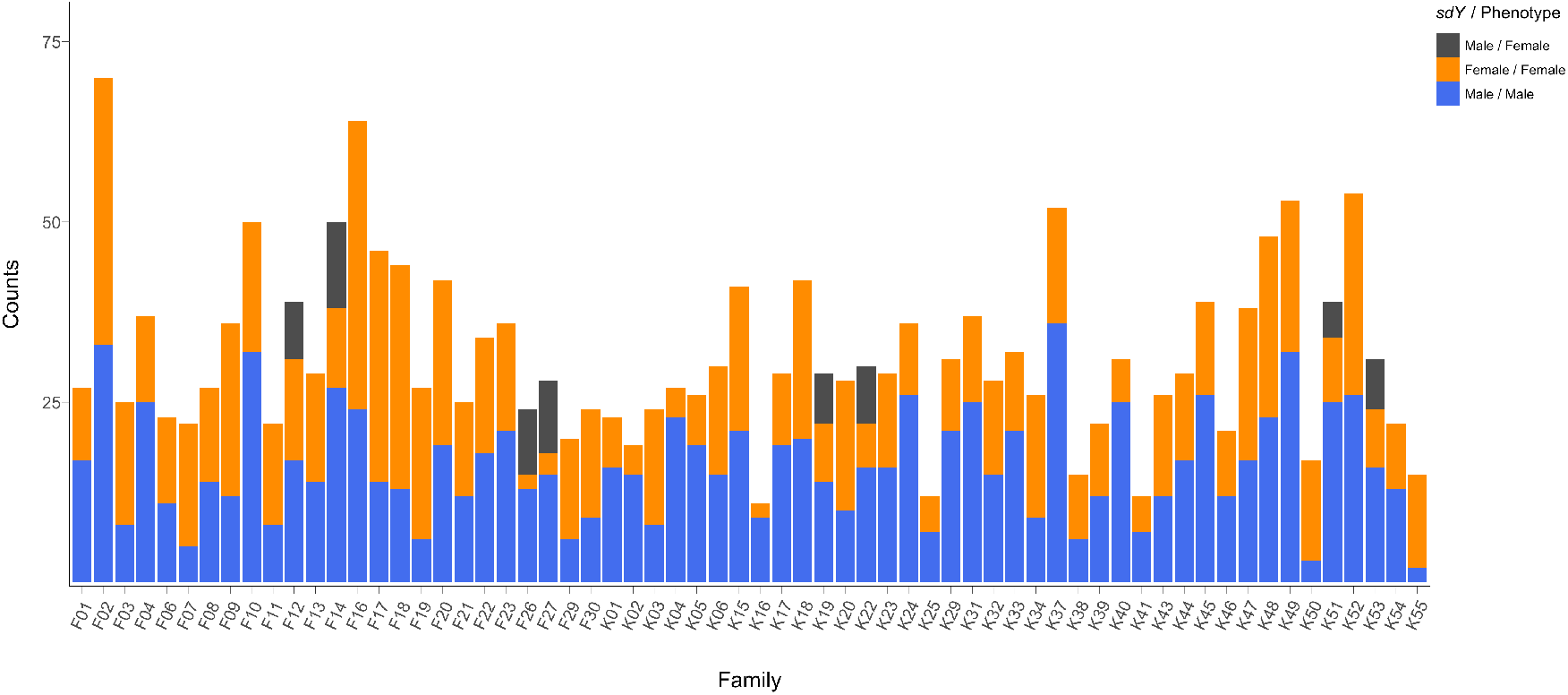
Phenotypic sex frequencies distribution across the 64 studied families showing PCR-based *sdY* genetic sex concordances. Concordant males and females are displayed in blue and orange respectively. Discordant females (*sdY* positive phenotypic females) are shown in dark grey.

Based on the above result, we hypothesized that Atlantic salmon may display a second autosomic copy of *sdY* in the genome. To examine this possibility, we used FC values from the qPCR assay in order to investigate the number of copies of the *sdY* present in each individual (both the sex determining gene and the potential autosomic pseudocopy). First, we genotyped known XY and YY males in order to validate the potential to identify two copies of this gene using the assay (Fig. 2a).

**Figure 2.**
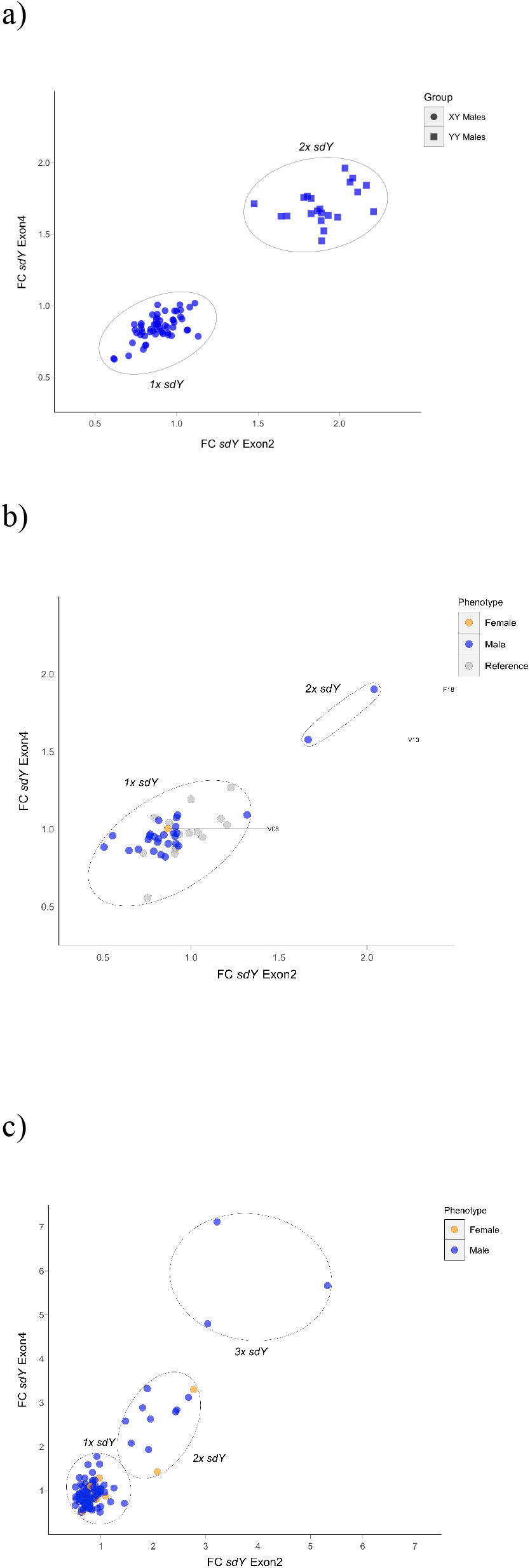
Real time PCR based fold change (FC) values for *sdY* exon2 and exon4 amplicons. (a) Validation RT PCR using XY males and YY super-males; round and square dots respectively. (b) Parental individuals for the F1-K2012 families showing 1-2x *sdY* sires individuals, in blue, and the 1x *sdY* discordant dam in orange. Reference 1x *sdY* males are displayed in grey(c) Example plate showing 1-3x *sdY* males and 1-2x *sdY* females in blue and orange respectively among the offspring. Reference males not displayed.

This test demonstrated that XY males clustered around 1 FC values for both amplicons (exon2 and 4). In addition, YY males, i.e., males containing two *sdY* copies, all clustered around a FC value of 2 (>1.5 FC threshold for both exons). All three discordant dams described above all carried a single *sdY* copy while they were four sires that carried two copies of *sdY* (Fig 2b). Significantly, these four males sired the five families displaying discordant offspring that did not have a discordant dam, and in addition, sired two families with a discordant dam (Table 1). Thus, at this stage, it was clear that all eight families displaying discordant offspring had one or two discordant parents. None of the 56 families without discordant offspring had discordant parents. Among the 2025 offspring, 62 and four of the phenotypic females displayed one and two copies of *sdY* respectively, and, 66 and eight phenotypic males displayed two and three copies of *sdY* respectively (Fig 2c). All of these individuals were reported from the eight families with offspring displaying discordance between phenotypic and genetic sex. When these data were considered on a family by family basis, the observed frequencies of the offspring carrying a variable number of copies of the *sdY* gene were highly consistent with autosomic mendelian inheritance (Fig. 3 and Fig. 4). Based upon our hypothesis, and the observed parental *sdY* genotypes in six of the families, we expected to see a 50/50 frequency in the female offspring displaying 0x vs. 1x *sdY* copies, and the same frequency in male offspring displaying 1x vs. 2x *sdY* copies. The observed frequencies (Fig. 3) did not conflict with the expected frequencies (Chi Square p values ranging from 0.24 to 0.98).

**Figure 3.**
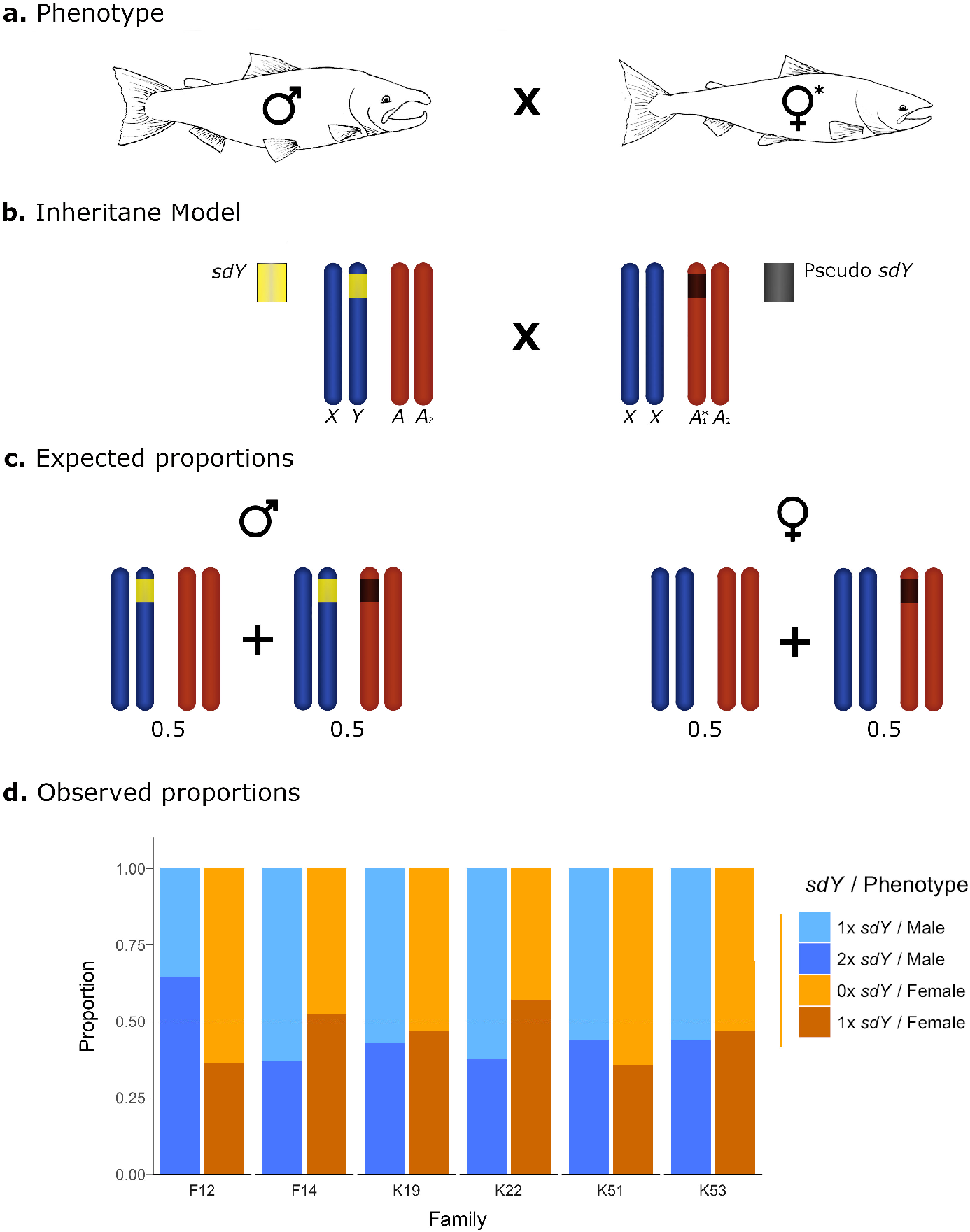
Conceptual diagram showing the inheritance model and the observed frequencies for the offspring of crosses with one parental individual carrying an *sdY* pseudocopy. (a) Cross between a 1x *sdY* phenotypic male and a 1x *sdY* discordant female denoted with *. (b) conceptual diagram showing the inheritance model at a chromosome level. Sex chromosomes and autosomes are represented in dark blue and dark orange respectively. Normal 1x *sdY* male (right) carrying a copy of the sex determining *sdY* gene (yellow) in the Y chromosome. Discrepant 1x *sdY* phenotypic female (left) carrying an *sdY* autosomic pseudocopy (dark grey) in heterozygosis. (c) Males and females expected proportions for the different *sdY* genotypes (1-2x). (d) Offspring observed proportions for the affected families of 1x and 2x *sdY* males and females: light blue, blue, light orange and orange respectively. Dotted lines represent the *sdY* genotype expected frequencies.

**Figure 4.**
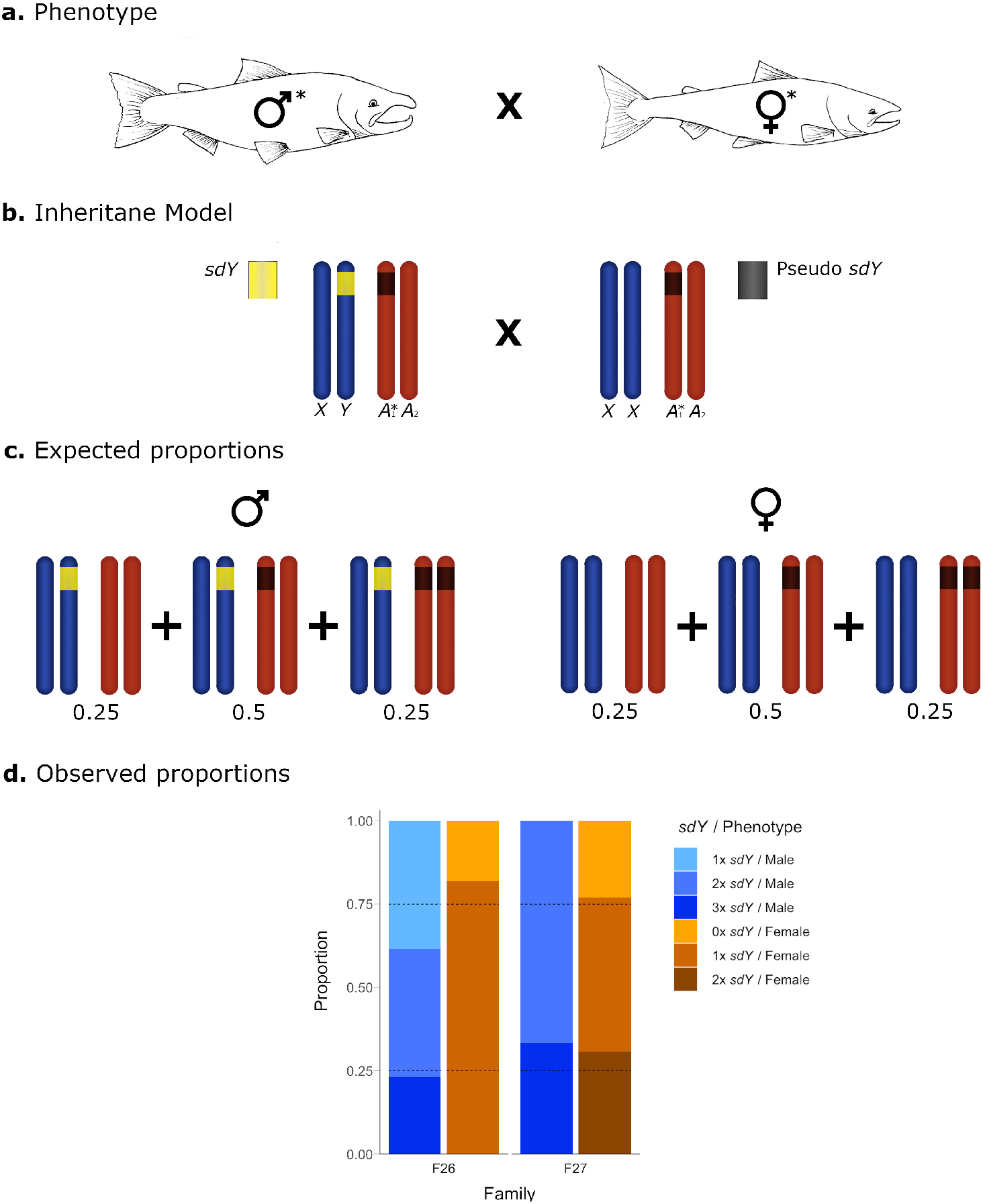
Conceptual diagram showing the inheritance model and the observed frequencies for the offspring of crosses with both parental individuals carrying an *sdY* pseudocopy. (a) Cross between a 2x *sdY* phenotypic male and a 1x *sdY* discordant female: Both affected parental individuals are denoted with an asterisk. (b) conceptual diagram showing the inheritance model at a chromosome level. Sex chromosomes and autosomes are represented in dark blue and dark orange respectively. (b) 2x *sdY* male (right) carrying a copy of the sex determining *sdY* gene (yellow) in the Y chromosome and the extra autosomic copy (dark grey) in heterozygosis. Discrepant 1x *sdY* phenotypic female (left) carrying an *sdY* autosomic pseudocopy (dark grey) in heterozygosis. (c) Males and females expected proportions for the different *sdY* genotypes (1-3x). (d) Offspring observed proportions for the affected families of 1x to 3x *sdY* males and 0 to 2x *sdY* females: light blue, blue, dark blue light orange, orange and dark orange respectively. Dotted lines represent the *sdY* genotype expected frequencies.

In the remaining two families, and based upon the parental genotypes, we expected to see a 25/50/25 distribution in the frequencies 0x, 1x, 2x, and 1x, 2x or 3x copies of *sdY* for female and male offspring respectively. Although the observed offspring frequencies did not match exactly with these expected frequencies (Fig. 4), most likely due to very low N offspring within these families, they did not significantly deviate from the expected frequencies (p values 0.29 and 0.34 from families 26 and 27 respectively.

Within the 64 families, validated phenotypic sex was mapped to chromosomes Ssa02, Ssa03 and Ssa06 (Table 1). Thereafter, the offspring from the eight affected families, with their *sdY* genotype (i.e., females displaying 0x vs 1x or 2x *sdY* copies, and males displaying 1x vs 2x, or 3x *sdY* copies), was mapped to chromosomes Ssa03, Ssa05, Ssa06, Ssa13 and Ssa28. Statistical support was however variable for some of the mapping data, in part possibly due to low N observations (Table 1).

## Discussion

The discovery of both the master sex-determining *sdY* gene and its function in salmonids represents a significant advance in knowledge (Yano et al. 2012; Yano et al. 2013; Bertho et al. 2018). However, phenotype-genotype discordances have been reported in many of the salmonid species, which has left open questions (Yano et al. 2012; Cavileer et al. 2015; Larson et al. 2016). Here, we have proposed the new hypothesis that discordance between phenotypic and genetic sex, at least in Atlantic salmon, is most likely caused by low-frequency *sdY* pseudocopies not involved in sex determination.

Genetic Sex determination and heteromorphic sex chromosome evolution is often tightly linked. Upon the arising of a novel sex-determining gene, sex chromosome evolution predicts the spread of reduced recombination between the sex-determination locus and linked genes with sex-specific fitness effects. Ultimately, reduced recombination will result in degeneration of sex-linked loci in the heterogametic sex and subsequent chromosome decay (Rice 1987; Charlesworth 2017). However, sex chromosomes may quickly evolve in many lineages and the sex chromosome pair may therefore change over time (Pennell, Mank, and Peichel 2018). This frequent sex chromosome turnover can provide a way to avoid chromosomal decay as has been shown in several fish species with young sex chromosomes (Kitano and Peichel 2012; Myosho et al. 2015).

In a species that has undergone recent whole genome duplications the possible existence of two or more copies of the sex determining gene can represent a major challenge (Ohno 1966). Duplicated sex determining genes can be removed, silenced or novel sex determining genes can be recruited. In Atlantic salmon, the presence of the Y specific *sdY* gene has been invoked to be the sole genetic requirement for maleness (Yano et al. 2013). Interestingly, the existence of multiple sex determining loci (Eisbrenner et al. 2014; Gabian et al. 2019), and *sdY* transposition ability (Lubieniecki et al. 2015) may give support to the existence of frequent sex chromosome turnover in this species. *sdY* male specificity is common in salmonids, but not in all species. Significantly, discordant genotypes i.e., *sdY* positive phenotypic females has been reported in Atlantic salmon (Eisbrenner et al. 2014) and other salmonid species (Cavileer et al. 2015; Larson et al. 2016). However, the existence of an inactive copy causing the apparent discordance has not been demonstrated until the present study.

Within the genetic material studied here, we observed 6.75% discordant females, and a total of 4% of the individuals with a second or third copy of the pseudo *sdY* gene. These originated from 3 dams and 5 sires among the 88 parents. The number of discordant females observed here is higher than the 1% frequency observed in a domesticated Tasmanian Atlantic salmon strain (Eisbrenner et al. 2014; Kijas et al. 2018). Given the inheritance model presented here, it is likely that this difference is merely the result of the number of affected parents and the cross design, although strain specific differences in the frequency of the pseudocopy of *sdY* cannot be ruled out. The 7% and 12% discordances between phenotypic and *sdY* determined sex reported in sockeye salmon (Larson et al. 2016) and the chinook salmon (Cavileer et al. 2015) respectively, raise two important questions. First, is the same mechanism demonstrated in Atlantic salmon here also the cause of observed discordance in the other salmonid species? Second, if the same mechanism is at play, is this generated as an example of parallel evolution or an event that happened early in the evolution of this genus? Members of the Coregoninae subfamily lack *sdY* male specificity and the existence of sex specific inactive copies in females has been invoked to explain this phenomena (Yano et al. 2012). *sdY* mobile nature may also explain the existence of inactive *sdY* in other salmonid species. By using the technical approach reported here, further research will shed some light into these important questions.

Within Atlantic salmon, *sdY* has been mapped to chromosomes Ssa02, Ssa03 and Ssa06 in a domesticated strain of salmon originating from North America, and, in both wild and domesticated Norwegian strains (Eisbrenner et al. 2014; Kijas et al. 2018; Besnier et al. 2020). It has also been mapped to chromosomes Ssa02 and Ssa21 in six wild Spanish populations (Gabian et al. 2019). The above observations are consistent with the findings of the present study where *sdY* was mapped to 2, 3 and 6 in the 64 families. In all these studies, chromosome 2 is identified as the most common location for *sdY* and is likely to be the ancestral variant (Kijas et al. 2018). Surprisingly, low divergence between the Ssa03 and Ssa06 loci has been reported (Kijas et al. 2018), suggesting a recent origin even though these variants are present in both the North American and European lineages. Convergent sex chromosome turnover might be the product of *sdY* transposable nature (Faber-Hammond, Phillips, and Brown 2015) and gene landscape (Bertho et al. 2018).

Here, we mapped the pseudo copy of *sdY* to chromosomes Ssa03, Ssa05, Ssa06, Ssa13 and Ssa28. Interestingly however, within the eight families displaying a pseudo copy of *sdY*, location of *sdY* and pseudo-*sdY* was only on the same chromosome in family 12. Therefore, these two genes typically do not co-locate on chromosomes. Given primer binding sequence conservation, these pseudocopies may represent recent transpositions and constitute the raw evolutionary material for sex chromosome turnover. Non-random transitions have been reported in amphibians (Jeffries et al. 2018) and seem to rely on finding the right genomic environment (Blaser, Neuenschwander, and Perrin 2014; Bertho et al. 2018). Particularly, *sdY* function relies on the spatial and temporal expression in some somatic cells of the early differentiating gonad, where inhibits female sex determination by interacting with key factors (Bertho et al., 2018). In medaka (Nanda et al., 2002), an autosomic locus is the origin of the sex determining gene and the former plays an active role in sex determination (Herpin et al., 2010). However, the *sdY* pseudocopies detected here may lack the regulatory regions needed for expression at the right time and place during sex determination. Furthermore, different chromosome pairs may be co-opted for sex in Atlantic salmon beyond Ssa02, Ssa03 and Ssa06. Kijas et al., (2018) found a significant association signal in Ssa25 and recently Ssa21 has been reported as candidate for sex in Spanish wild populations suggesting an alternative phylogeographic origin. Besides the domesticated and wild populations analyzed here, we have found *sdY* pseudocopies in wild salmon from the river Årdalselven in Norway (data not presented). It is then, plausible, if not likely, that this is a common phenomenon in wild populations. However, the geographic, biological and life-history patterns of this need further research. For this purpose, our method to quantify genomic copies of the *sdY* yields an accurate estimate of the frequency of these discordances.

Results of the present study, including the *sdY* copy number assay developed herein, have implications for commercial salmonid breeding programs. Breeders are increasingly using *sdY* to determine phenotypic sex and to assist broodstock selection in the early production phase. Discordance between phenotypic and *sdY* based genetic sex has been reported for example in the commercial Mowi strain (Matt Baranski, Mowi, personal communication). This represents a logistic and financial challenge as maturing males and females are transferred to land-based rearing facilities and held separate until spawning. Mis-identified females take precious space and seriously impact cross design later on. Being able to identify both males and females carrying pseudocopies will allow to remove this pseudocopy out of the breeding line in one generation: double/triple males and double females can be removed early, and single females weeded out when the phenotype is clear. Additionally, gaining knowledge about the proper genomic environment needed for a successful sex chromosome turnover might significant a huge leap in the race of understanding the precise mechanisms behind sex determination and ultimately in gaining control of the process from an aquaculture perspective.

## Acknowledgements

This work was funded by the Norwegian Ministry of Trade and Fisheries, and the Norwegian Research Council in the grants INTERACT (200510) and MATGEN (254783). Neither funding body played any role in the design of the study, interpretation of data, nor conclusions drawn. We greatly acknowledge Lise Dyrhovden, Ivar Helge Matre, Kåre Storsæter and Jan Olav Fosse at the Matre research station (IMR) for rearing of fish included in this study, Anne Grete E. Sørvik and Zhiwei Zhang for microsatellite and SNP genotyping, and MOWI for unconditionally providing the domesticated gametes and for discussions regarding the implications of these results. We would also like to thank the river owners for access to wild broodstock. Emily K. Glover is gratefully acknowledged for drawing the salmon used in Figures 3 and 4.

---

**Supp. Table 1.**
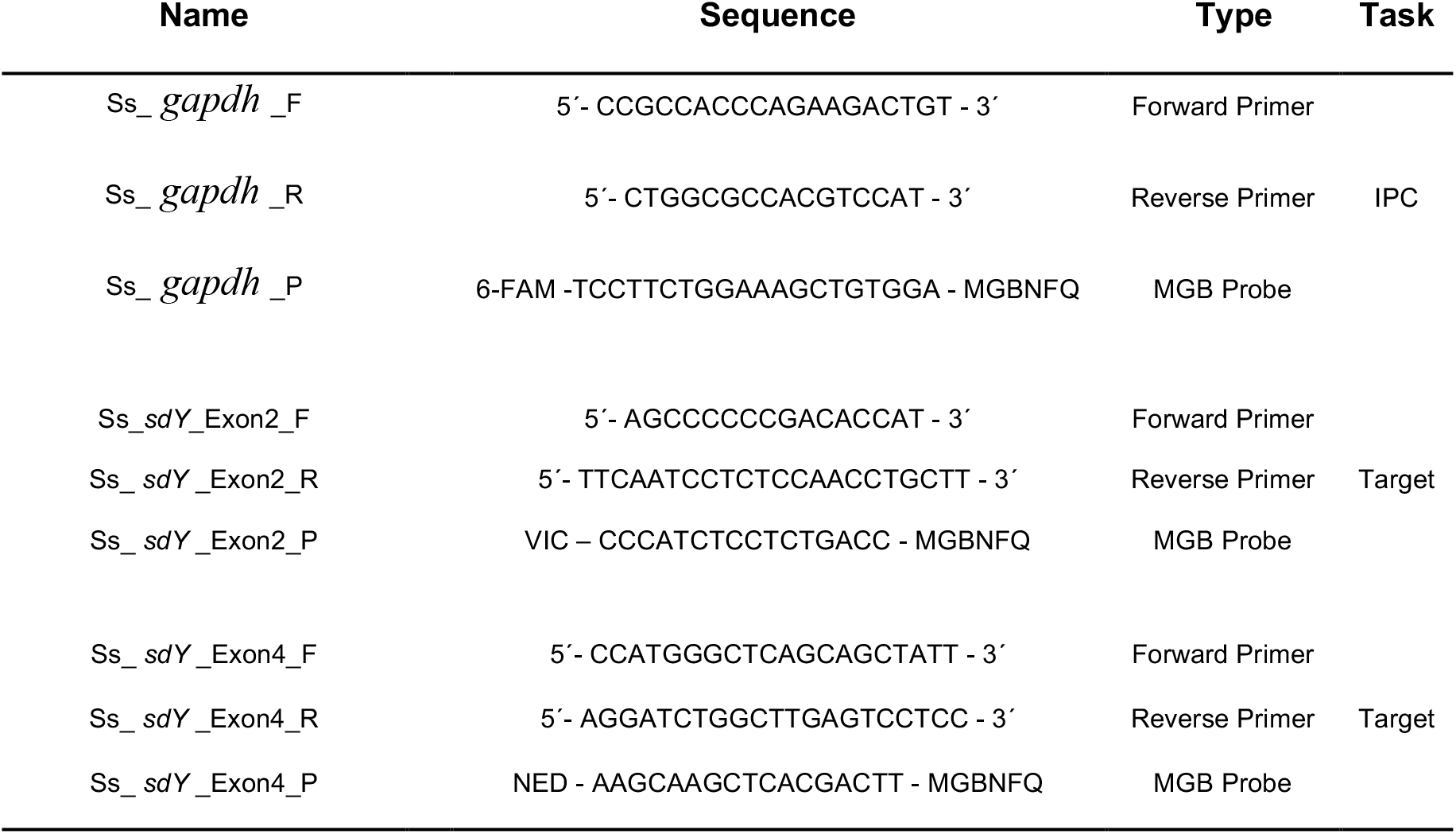
Primers and custom 5’ labeled probes used for real time PCR genetic sex determination. 6-Fam, VIC and NED reported dyes were used for *gapdh*, sdY exon 2 and 4 respectively. *gapdh* was used as internal positive control (IPC) and reference for fold change calculations.

